# Gastric administration of C*is*-9, *trans*-11 and *trans*-10, *cis*-12 conjugated linoleic during the pregestational and gestational periods does not influence the follicular endowment of the progeny

**DOI:** 10.1101/2022.10.06.511095

**Authors:** Danielle Storino de Freitas, Guilherme Antonio de Gouvêa Lopes, Barbara Rodrigues Nascimento, Ana Paula Madureira, Paulo Henrique Almeida Campos-Junior

## Abstract

Fetal programming suggests that maternal stimulation and nutrition during the period of fetal development can program the progeny. Conjugated linoleic acid (CLA), an isomer of linoleic acid, has been characterized in several aspects, but few studies have been performed on its involvement in reproduction and fetal programming. The aim of this study was to evaluate the F1, F2 and F3 progeny of female mice supplemented with CLA during the pregestational and gestational periods with respect to biometric and reproductive parameters, as well as ovarian morphophysiology. The F1 progeny of mothers supplemented with CLA exhibited stable weight gain, while the F2 progeny showed no effects (P=0.0187 and P=0.0245, respectively). A reduction in Lee’s Index was observed in both generations at the second post-weaning evaluation week in the animals treated with CLA (P=0.0100 and P=0.0078, respectively). The F2 generation showed an increase in the anogenital index in both sexes of the animals treated with CLA (P= 0.0114 and P<0.0001, female and male respectively). CLA administration to mothers did not affect any of the following in their progeny: ovarian follicle mobilization (P>0.05), follicle number (P>0.05) and the integrated density of the lipid content of oocytes included in antral follicles (P>0.05). This study evaluated the use of CLA in mothers and found that it did not affect the progeny regarding murine reproductive performance, suggesting that this supplement can be used safely.

## Introduction

Historically, studies have been conducted to understand how maternal nutrition influences the development and offspring health [1]. The concept of fetal programming suggests that maternal stimulation and nutrition during the period of fetal development can program the progeny into adulthood [2,3].

Nutritional demands and their effects are considered individual, as is the response of offspring to maternal nutrition factors. The effects of maternal nutrition during pregnancy on the offspring’s health may be more apparent during the perinatal period [4]. The fetus adaptation to the morphological, physiological and molecular pressures submitted during the embryonic development are made possible by epigenetics [5], characterized by molecular factors or processes around DNA that have the ability to regulate genomic activity independently of the DNA sequence and are both mitotically stable and heritable through the germ line [6].

Some genes can remain imprinted in specific tissues of the organism throughout life [7], however, with DNA methylation, the changes can be transmitted transgenerationally [8]. Thus, these processes are extremely important in the regulation of genomic activity [9] in animals exposed to numerous factors during gestational development. Conjugated linoleic acid (CLA) refers to a mixture of positional and geometric isomers of linoleic acid with conjugated double bonds, not separated by a methylene group as in linoleic acid. These isomers can be synthesized in the rumen, adipose tissue and ruminant mammary gland, a process known as endogenous synthesis. CLA is widely used as a food supplement, due to its ability to maximize the use of body fat reserves, reduce carcinogenesis, exert an obesity effect, and alter the lipid composition of bovine milk [10]. The supplementation of diets and culture media with CLA is an emerging area of research, and studies are needed to elucidate its beneficial effects on reproductive parameters. Recently, we demonstrated that gastric administration of CLA during the pregestational and gestational periods did not affect ovarian follicle endowment and mobilization, or did it affect oocyte lipid accumulation, demonstrating that this supplement can be used to take advantage of the benefits described in the literature without detrimental effects on female reproductive healthy mice [11]. Therefore, the aim of this study was to evaluate the effects of maternal gastric administration of CLA on the biometric parameters, ovarian morphophysiology and the fertility of the F1, F2 and F3 progeny in mice.

## Materials and Methods

### Animals, Facilities and Experimental Design

Female (n=30) and male (n=15) mice C57BL/6 were obtained from the Núcleo de Criação de Animais de Laboratório and maintained in an environment with a controlled temperature of 22 ±2 °C and with artificial light cycles (12:12 h). The 6 weeks old females were randomly selected and distributed in three groups: (1) control (n = 10), (2) fish oil (n = 10), and (3) CLA (n = 10), that daily received by gavage 35 μl of phosphate buffer saline (PBS), fish oil (Mundo dos Óleos^®^, Brasilia, DF, Brazil), synthetic CLA (Tonalin TG80 Basf^®^, São Paulo, SP, Brazil; 80% C18:2 conjugated, 39.2% C18:2 *cis*-9, *trans*-11 and 38.4% C18:2 *trans*-10, *cis*-12), respectively. The fish oil administration was used as a positive control, once the structural and functional similarities [12] with the fatty acid CLA profile is well established, and also there is a report about its effects on ovulation rate, and litter size in mice [13]. Treatments were performed during 50 days (before mating, mating and pregnancy) [11]. These females were crossbred with 10 weeks old untreated males (1:2, male:female) with proven fertility for a period of 2 weeks. Pups from the parental generation was called the F1 generation. F1 females were mated with untreated male mice at 6 weeks old to produce the F2 generation. F2 females were mated with untreated male mice at 6 weeks old to produce the F3 generation. After parturition, females were euthanized by CO_2_ asphyxia and their ovaries were collected. All experiments were conducted according to the principles and procedures described by the Brazilian National Board of Animal Experimentation Control (CONCEA) and were approved by the Institutional Ethics Committee of the Federal University of São João del-Rei (Protocol 013/2018).

### Biometric data

The animals were weighed weekly on a commercial scale (SF-400^®^, São Paulo, SP, Brazil), and nasoanal length also was measured. The Lee index (obesity indicator) was calculated as previously described by [14]. The anogenital length was measured at the 2nd day old, calculated according to the formula proposed by Whelsh [15], considering IAG=DAG/weight (mm/kg).

### Ovarian follicular quantification

All ovaries were fixed in PFA (Sigma-Aldrich, St. Louis, MO, USA) solution. Paraffin-embedded ovaries were serially sectioned (5 μm) and stained with hematoxylin and eosin solution. In every fifty sections, the number of primordial, primary, secondary, antral, and atretic follicles were quantified, estimating the total number of follicles per ovary. Follicles were classified and counted [16] and follicles (from primordial to antral) showing morphological signs of death such as pyknosis, cellular fragmentation, and disintegration were classified as atretic [16]. Only follicles containing an oocyte with a visible nucleus were considered to avoid double-counting. Results are shown as the number of counted follicles per animal. Atresia rate (%) was determined as the number of atretic follicles/total number of follicles*100; and activation rate (%) as the number of growing follicles/ total number of follicles*100 [16].

### Morphometric evaluation

Follicle diameters were calculated from the average of two perpendiculars, using the image analysis program Software ImageJ (NIH, USA). The follicle boundary was defined with the basement membrane clearly visible, as a demarcation between the granulosa cells and a special theca, and the oocyte boundary was the zona pellucida. Thirty follicles of the class (primary, secondary and antral) were randomly selected for treatments. Only follicles completely free of any signs of atresia were analyzed.

### Oocyte Lipid Quantification

Paraffin-embedded ovaries were also serially sectioned (5 μm) and stained with Sudan red dye, as described by Sudano [17]. The Sudan red dye solution was prepared with 3% Sudan IV in 70% ethanol. To estimate the relative amount of lipid droplets per oocyte, images of antral follicles (12 oocytes per animal; 6 animals per group) were captured in 40X objectives. All photos were converted to black and white (32 bt), and the oocytes included in antral follicles were delimited and analyzed using the area of integrated density compared. This evaluation was performed in the Image J^®^ software according to Leite [18].

### Statistical analysis

Statistical analysis was carried out using statistical software (GraphPad Prism 8.0, Graph Prism Inc., San Diego, CA). A completely randomized design (DIC) was carried out with the following mathematical model:

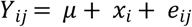

Where,

*Y_ij_* = Bodyweight, nasoanal length, Lee index, primordial, primary, secondary, antral, total and atretic follicles, atresia and activation rate
μ = Constant
*x_i_* = Supplementation (Control, fish oil, CLA)
*e_ij_* = Random error

For the characteristics bodyweight, nasoanal length and Lee index, an intercepts and slope comparison of the line was performed by means of a simple linear regression analysis during the 4 weeks of evaluation. For the variables number of primordial, primary, secondary, antral, total and atretic follicles, atresia and activation rate a D’Agostino & Pearson normality test was conducted to test the null hypothesis that the data are sampled from a Gaussian distribution. When the data did not deviate from Gaussian distribution (P>0.05) they were analyzed using the one-way ANOVA with Newman-keuls test for multiple comparisons. Otherwise, a Kruskal-Wallis test was done with a Dunn’s test for multiple comparisons. A Student t-test was applied to Body Weight, NA and Lee Index in two-group comparisons. All tests were performed at a level of significance of 0.05.

## Results

In the F1 generation, the treatments did not affect body weight in the first week of post-weaning evaluation, however, a difference was observed in the second and third weeks (P=0.0187 and P=0.0245, respectively) in which the weight of animals in CLA treatment increased in comparison to the others, stabilizing in the fourth week (Table 1). In F2 generation, exposure of the animals to CLA showed similar results to those in the control over the four weeks of evaluation (P=0.0005, P=0.0034, P=0.0212 and P=0.0561, respectively) (Table 1).

**Table 1.** Effects of prenatal exposure to control, fish oil and CLA on body weight, measured in milligrams, in the F1 and F2 generations of female mices.

| Time point | Treatments | Generation |  |  |  |
| --- | --- | --- | --- | --- | --- |
|  |  | F1 |  | F2 |  |
| 5 Weeks Post-Natal | Control | 15.36 ± 1.5 |  | 15.8 ± 1.9 | ab |
|  | Fish Oil | 15.3 ± 1.8 |  | 17.5 ± 0.5 | a |
|  | CLA | 16.4 ± 1.4 |  | 12.0 ± 2.5 | b |
| 6 Weeks Post-Natal | Control | 16.4 ± 1.3 | a | 17.4 ± 2.8 | ab |
|  | Fish Oil | 16.6 ± 1.6 | a | 20.0 ± 1.6 | a |
|  | CLA | 17.8 ± 0.9 | b | 14.3 ± 2.4 | b |
| 7 Weeks Post-Natal | Control | 17.9 ± 1.8 | ab | 19.6 ± 3.2 | ab |
|  | Fish Oil | 17.3 ± 1.8 | a | 20.5 ± 1.7 | a |
|  | CLA | 19.3 ± 1.9 | b | 15.4 ± 2.8 | b |
| 8 Weeks Post-Natal | Control | 19.7 ± 1.9 |  | 24.8 ± 2.8 | ab |
|  | Fish Oil | 18.9 ± 1.3 |  | 20.0 ± 1.0 | a |
|  | CLA | 20.7 ± 2.4 |  | 16.5 ± 2.3 | b |

Table represent means ± standard error of the mean. $P \leq 0.05$ (significant difference compared to the control)

A reduction in the Lee Index was observed in both generations in the second week of post-weaning evaluation (P=0.0100 and P=0.0078, respectively) of CLA-treated animals (Table 2).

**Table 2.**
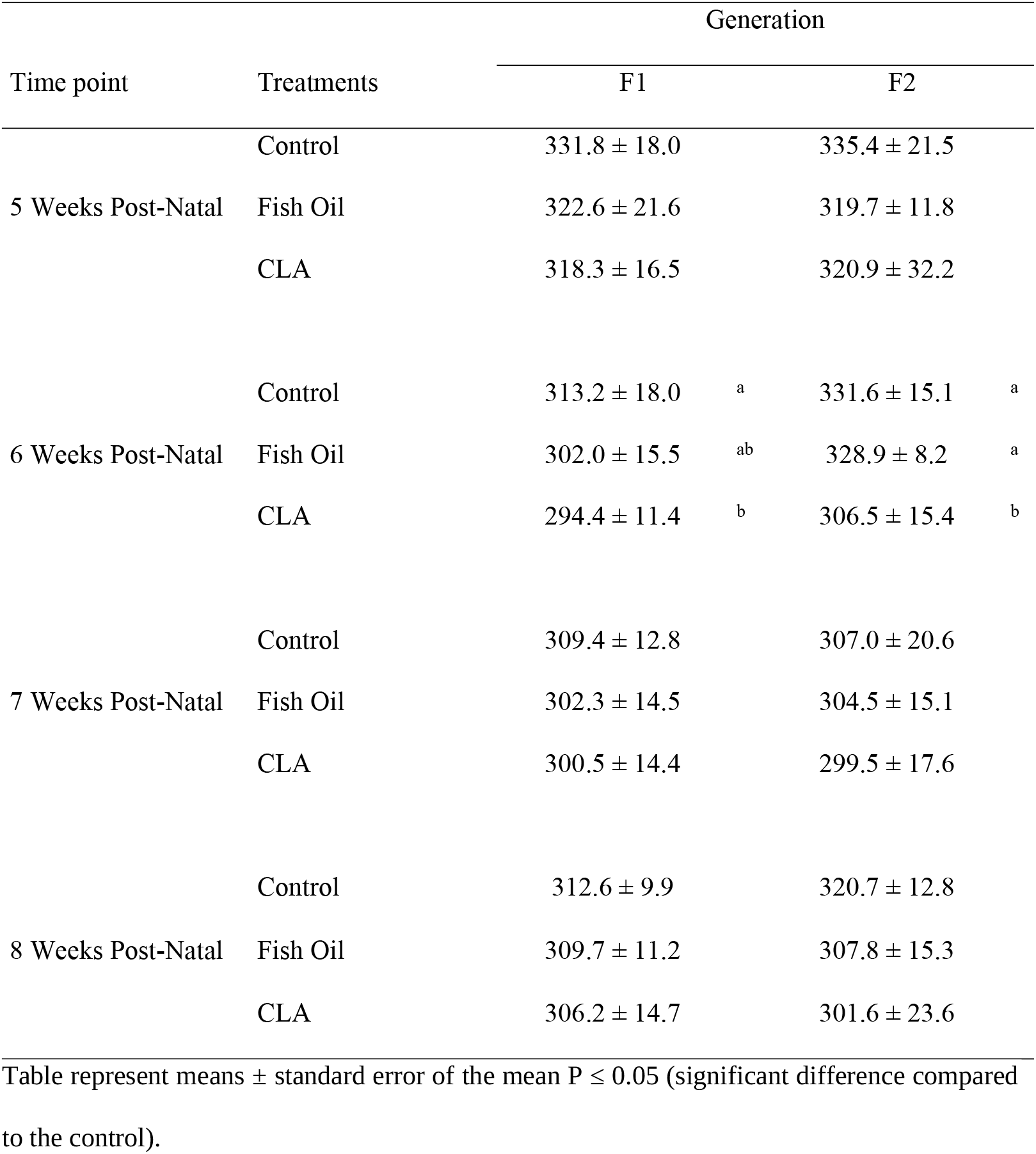
Effects of prenatal exposure to control, fish oil and CLA on Lee index in the F1 and F2 generations of female mices.

| Time point | Treatments | Generation |  |  |  |
| --- | --- | --- | --- | --- | --- |
|  |  | F1 |  | F2 |  |
| 5 Weeks Post-Natal | Control | 331.8 ± 18.0 | | 335.4 ± 21.5 | |
| | Fish Oil | 322.6 ± 21.6 | | 319.7 ± 11.8 | |
| | CLA | 318.3 ± 16.5 | | 320.9 ± 32.2 | |
| 6 Weeks Post-Natal | Control | 313.2 ± 18.0 | a | 331.6 ± 15.1 | a |
| | Fish Oil | 302.0 ± 15.5 | ab | 328.9 ± 8.2 | a |
| | CLA | 294.4 ± 11.4 | b | 306.5 ± 15.4 | b |
| 7 Weeks Post-Natal | Control | 309.4 ± 12.8 | | 307.0 ± 20.6 | |
| | Fish Oil | 302.3 ± 14.5 | | 304.5 ± 15.1 | |
| | CLA | 300.5 ± 14.4 | | 299.5 ± 17.6 | |
| 8 Weeks Post-Natal | Control | 312.6 ± 9.9 | | 320.7 ± 12.8 | |
| | Fish Oil | 309.7 ± 11.2 | | 307.8 ± 15.3 | |
| | CLA | 306.2 ± 14.7 | | 301.6 ± 23.6 | |

Table represent means ± standard error of the mean $P \leq 0.05$ (significant difference compared to the control).

In the F1 generation, the treatments did not affect the anogenital index in males or females (P>0.05) (Table 3). However, the F2 generation showed an increase in the anogenital index in both sexes of CLA-treated animals (P=0.0114 and P<0.0001, female and male respectively) (Table 3). and the male offspring of the F2 generation (F3 puppies) showed differences compared to the control and fish oil groups (P<0.0001) (Table 3).

**Table 3.** Effects of prenatal exposure to control, fish oil and CLA on anogenital index in the F1 and F2 generations of female mices.

| Treatments | Generation |  |  |  |  |  |
| --- | --- | --- | --- | --- | --- | --- |
|  | F1 |  | F2 |  | F3 |  |
|  | Female | Male | Female | Male | Female | Male |
| Control | 613.4 ± 54.8 | 1114.0 ± 201.2 | 588.7 ± 100.9<sup>a</sup> | 971.3 ± 136.0<sup>a</sup> | 488.9 ± 66.4<sup>a</sup> | 850.0 ± 54.2<sup>a</sup> |
| Fish Oil | 516.8 ± 84.4 | 968.8 ± 175.1 | 560.8 ± 94.44<sup>a</sup> | 965.0 ± 171.7<sup>a</sup> | 519.0 ± 79.0<sup>a</sup> | 1151.0 ± 71.94<sup>b</sup> |
| CLA | 618.7 ± 136.5 | 1010.0 ± 165.4 | 690.7 ± 139.5<sup>b</sup> | 1130.0 ± 134.6<sup>b</sup> | 578.8 ± 119.5<sup>b</sup> | 1037.0 ± 106.0<sup>c</sup> |

Table represent means ± standard error of the mean $P \leq 0.05$ (significant difference compared to the control)

Regarding morpho-quantitative analyses, there was no difference (P>0.05) in the number of primordial, primary, secondary, antral, atretic, and total follicles per ovary in control, fish oil, and CLA-treated animals in the F1 and F2 generations (Fig 1 and 2).

**Fig 1.**
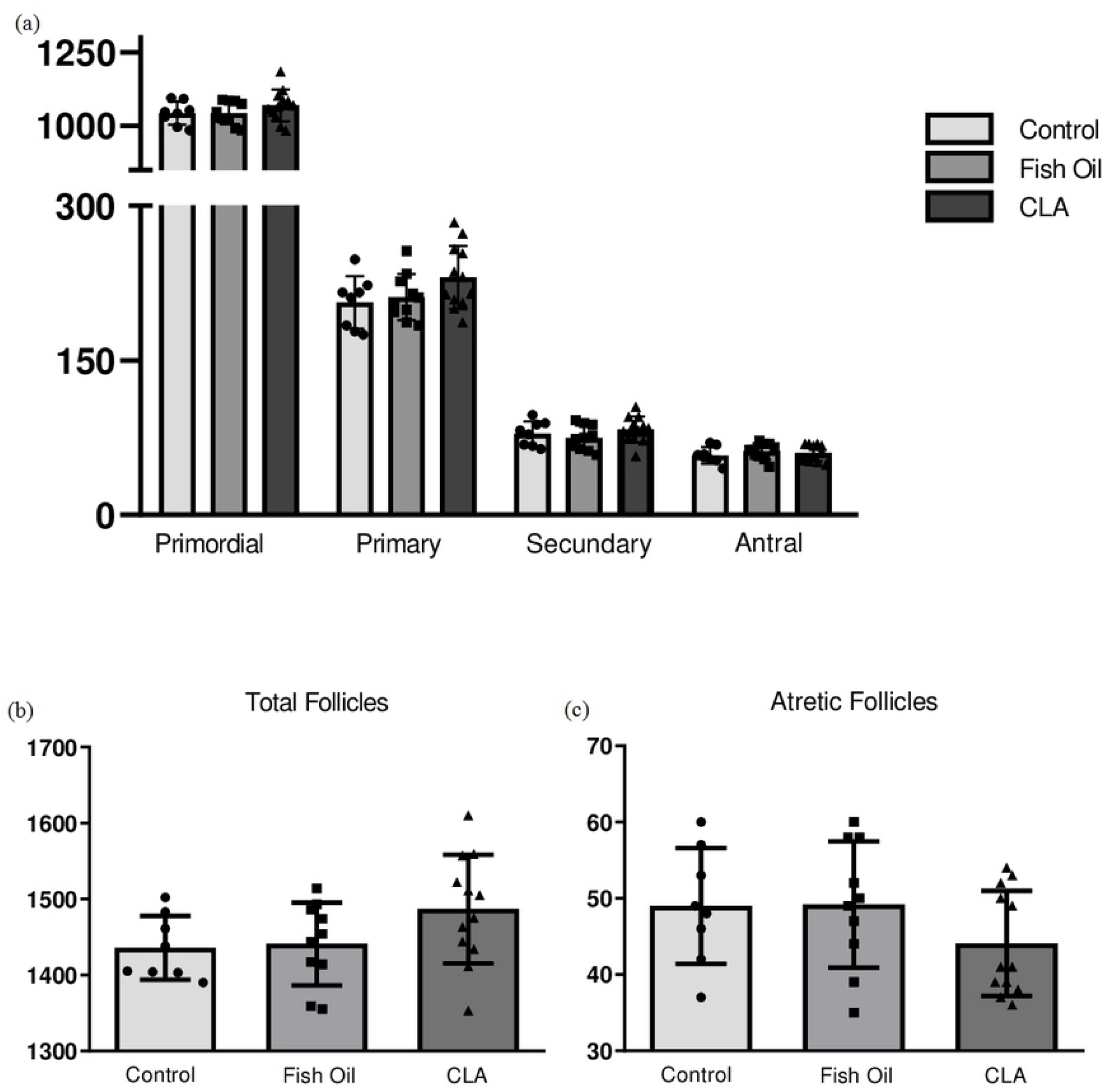
(a) Primordial, primary, secondary and antral follicles, (b) total follicles and (c) atretic follicles of ovaries from control, fish and CLA of F1 generation. These parameters were not altered by treatment (P>0.05).

**Fig 2.**
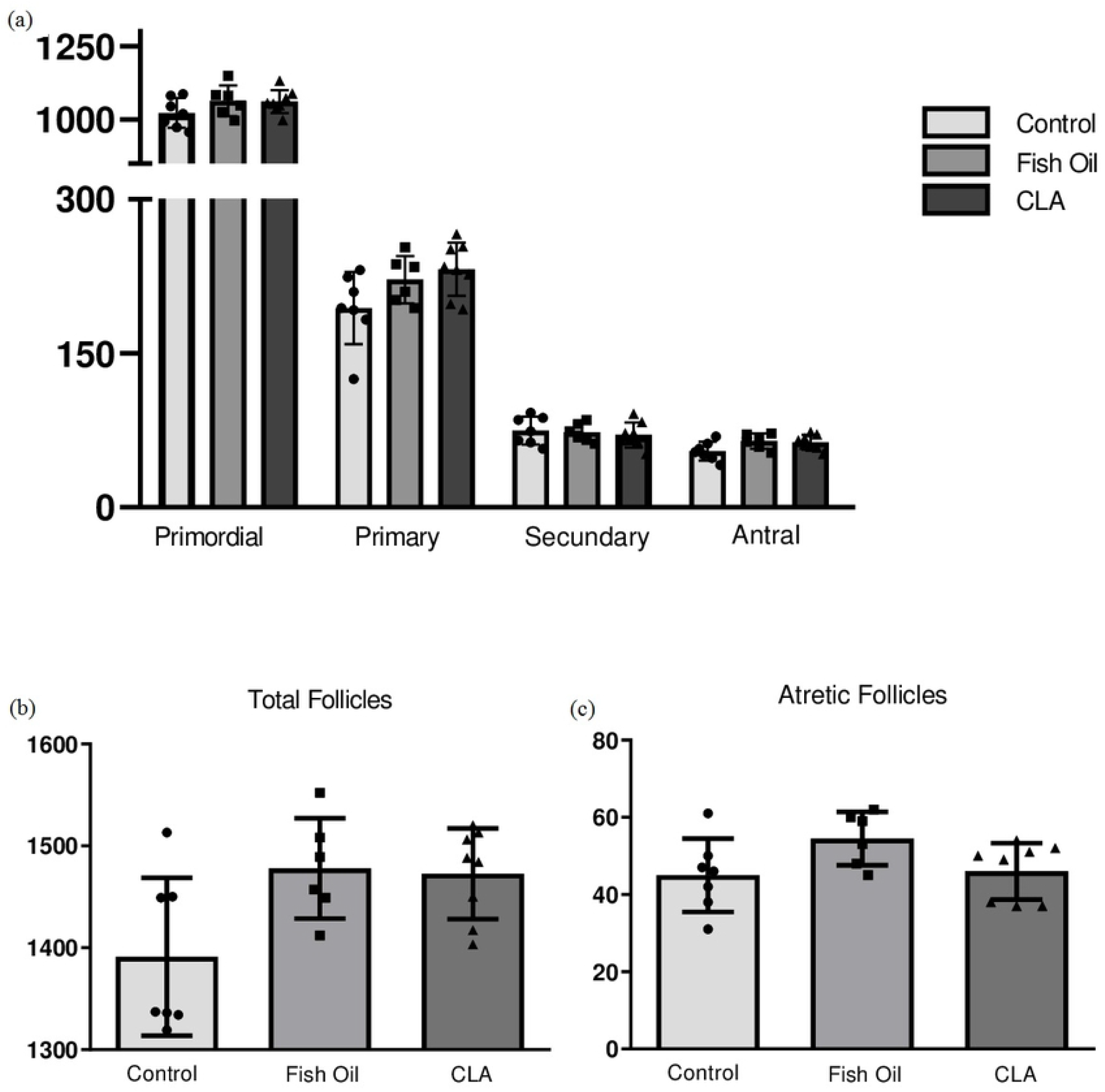
(a) Primordial, primary, secondary and antral follicles, (b) total follicles and (c) atretic follicles of ovaries from control, fish and CLA of F2 generation. These parameters were not altered by treatment (P>0.05).

All classes of follicles were observed qualitatively observed in the ovaries of control, fish oil, and CLA treated animals and no morphological alterations resulted from these treatments over two generations (Fig 3), as the follicle and oocyte diameter were not altered (P>0.05), nor was the oocyte:follicle ratio (P>0.05).

**Fig 3.**
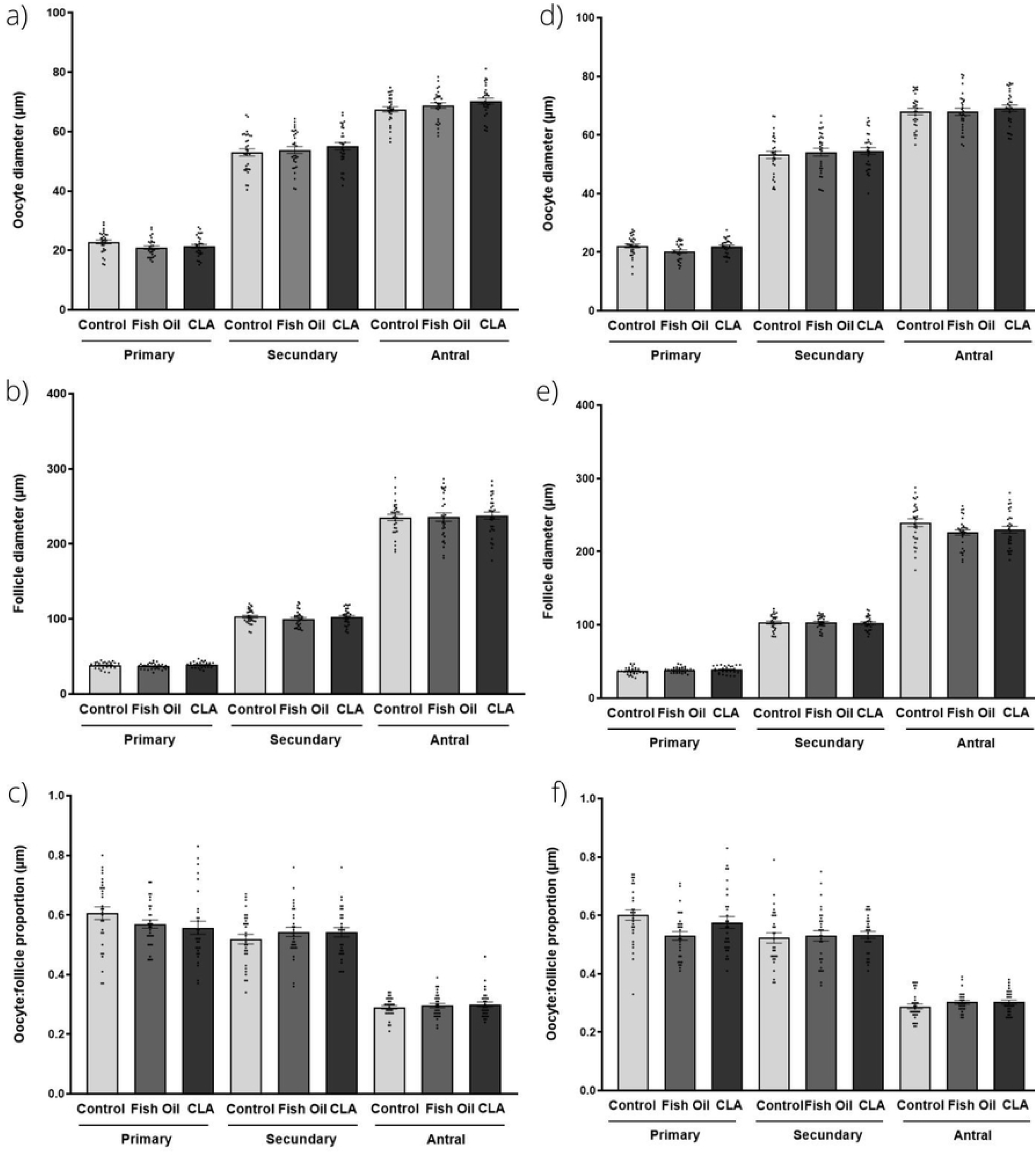
Morphometry of primary, secondary and antral follicles from control, fish oil and CLA groups. (a) Diameter of the oocyte; (b) follicle (c) oocyte:follicle diameter ratio of F1 generation, (d) diameter of the oocyte; (e) follicle and (f) oocyte:follicle diameter ratio of F2 generation. These parameters were not altered by treatment (P>0.05).

Additionally, an increase in the amount of lipid droplets, measured by Sudan IV staining, was observed in F1 animals from the CLA group compared to the fish oil group (P=0.0446) (Fig 4A). The F2 generation showed no difference in this parameter (P>0.05) (Fig 4B).

**Fig 4.**
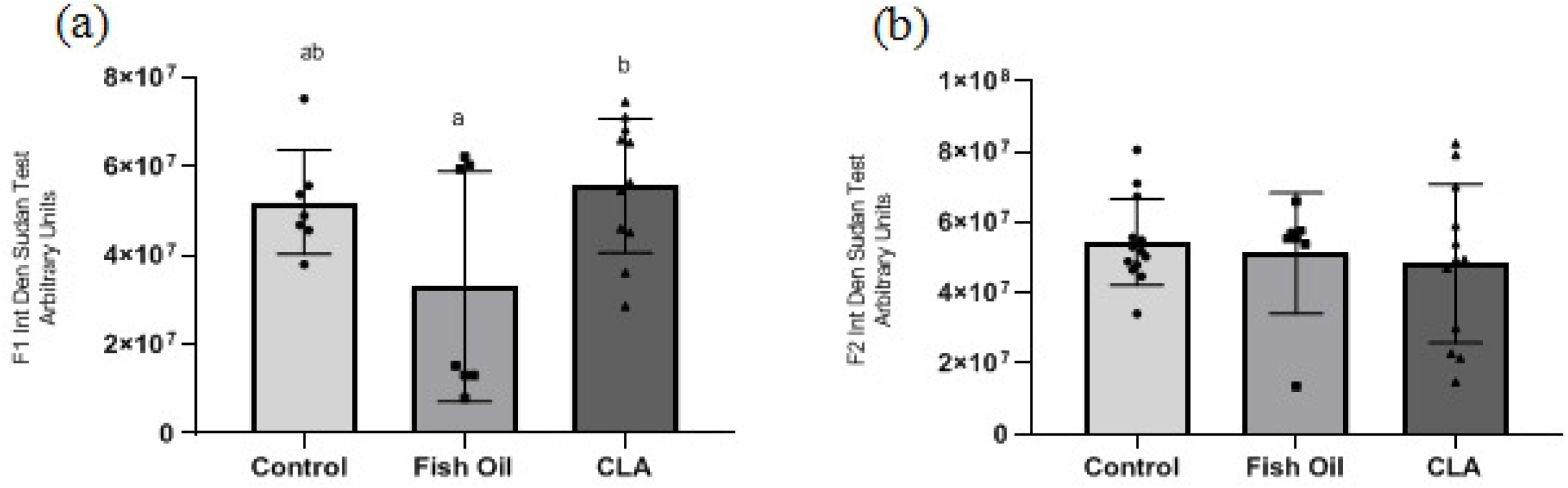
Integrated density of lipid content (arbitrary units) of oocytes included in antral follicles among control, fish oil, and CLA groups for F1 (a) and F2 (b) generation.

## Discussion

Organogenesis and tissue differentiation occur during the prenatal period, considered a critical moment for the programming of the offspring’s phenotype, which can impact on the postnatal period, persisting into the offspring’s adult life [19]. Studies carried out in rodents demonstrated the nutritional effects on GnRH production and secretion, which can affect female puberty [20]. CLA supplementation has demonstrated some beneficial effects, such as the regulation of adipogenesis and myogenesis and alteration of body composition [21], as well as immunological, cardiovascular and anticarcinogenic activities [22, 10]. However, few studies have evaluated the reproductive parameters of the offspring of CLA-treated females [13]. Thus, our data demonstrated that the use of CLA through gastric administration in the parental generation did not affect the obesity rates and follicular mobilization in later generations, affirming the absence of side effects of gastric CLA administration on offspring.

The current study demonstrated that animals born to mothers supplemented with CLA did not show alterations in body weight and the obesity index, as previously reported for parental [11] and corroborating the literature [23]. Some studies evaluated weight gain in CLA-treated animals and reported similar findings. According to Peng [24], the *trans*-10, *cis*-12 isomer can pass through the umbilical cord from mothers to their progeny. Thus, the anti-obesity effects can be long-lasting. Studies carried out in dairy cows showed that the inclusion of CLA in the diet did not affect food consumption [25, 26], which may be lower in the maintenance of the animals weight throughout the life cycle time. Although the potential of CLA to AID in weight loss is publicized, Vaisar et al. [27] and Gaullier et al. [28] indicate that in healthy individuals, this loss is less pronounced than in those who already have obesity.

In mammals, the puberty onset is a physiological process marked by increased levels of steroid hormones, during which animal and humans attain the characteristics of an adults, such as throught sexual maturation and the development of reproductive and neurological organs [29]. Although such events are genetically determined, several factors can alter the outcome, such as nutritional and metabolic status, as well as environmental factors [30]. For the beginning of this process, the secretion of GnRH by the hypothalamus, which stimulates sex hormones through the pituitary and gametogenesis, is necessary, but the trigger for this stage of development has not yet been fully elucidated [29].

The sexually dysmorphic anogenital index in rodents is considered a broad biomarker of androgen exposure during the fetal masculinization programming window and predictor of late reproductive disorders in offspring [31, 32], however, the reflex of androgen action occurs within a reduced time interval in rats (days 15.5 to 18.5) [33]. The observed effect of CLA on the increase in the anogenital index in male and female mice can be attributed to the possible effect of this supplementation acting on the increase in the production of steroid hormones and may also be due to the exposure to androgens. Studies have demonstrated that neonatal exposure to androgens in women results in virilization by this exogenous pathway, both before and during the masculinization programming window [34]. This index in females, as shown in studies by Mendiola et al. [35] suggests that the androgenic environment during the prenatal phase may indicate an increase in the number of ovarian follicles and high levels of testosterone. However, our findings do not show variations in the number of follicles in the treatments evaluated.

In females, a greater anogenital distance can lead to a hyperandrogenic uterine environment, which may be associated with prenatal ovarian dysfunction [34]. While its shortening in males was associated with worse semen quality and the possibility of infertility [34]. Thus, it is possible that CLA has androgenic effects that lead to differences in the anogenital distance between males and females from generations that were exposed to CLA during fetal development. According to Bach [36], unsaturated fatty acids are direct substrates for the cholesterol production and, consequently, of steroid hormones, with an increase in the progesterone production, which is related to increased fertility and embryonic development [37].

Freitas et al. [11] found no effect on mobilization and follicular morphology, similarly to the present study. Campos-Junior et al. [16] state that several factors can act on these parameters: the inclusion of fat in the diet has been reported to increase follicle number and diameter [38, 39]. Moreover studies claim that the content of fatty acids in the follicular and oocyte fluid influences the development competence [40]. Broughton et al. [41] observed that the consumption of CLA did not influence the ovulation rate or the production of prostaglandins, inferring the difficulty of predicting the change through the inclusion of fatty acids. Csilik et al. [38] determined through studies with dairy cows that the best time to administer CLA is before calving, in which its accumulation would be advantageous to facing metabolic challenges. Our findings corroborates those of Yi et al. [13], who demonstrated that CLA supplementation did not affect the ovulation rate in mice.

Oocytes can accommodate large amounts of lipid droplets, and their absorption in the embryos is critical, being the main factor responsible for the accumulation of ATP through mitochondrial oxidation [42]. Abazarikia et al. [43] state that the inclusion of CLA in oocyte maturation and in vitro embryo culture can affect the modulation of lipogenesis. The data in the current study corroborate those of Freitas et al. [11], as supplementation of the maternal diet CLA, did not alter the lipid content of oocytes included in antral follicles.

In conclusion, our study showed for the first time in the literature that gastric CLA administration during the pregestational and gestational periods did not affect ovarian follicle endowment and mobilization in the F1 and F2 progeny. However, some punctual alterations in the anogenital and Lee indexes, as well as inbody weight, resulted from this treatment. Therefore, these findings indicate that CLA supplementation does not have a significant adverse effect on the parameters evaluated herein and can be used without apparent detrimental effects on female reproductive health on mice, taking advantage of the other benefits already described in the literature.

## Acknowledgements

This work was supported by the National Council for Scientific and Technological Development (CNPq, Brazil) and Minas Gerais State Research Foundation (FAPEMIG, Brazil). DFS, GAGL and BAM received a scholarship from CNPq and CAPES.

## Funding

This work was supported by the National Council for Scientific and Technological Development (CNPq, Brazil) and Minas Gerais State Research Foundation (FAPEMIG, Brazil). DFS, GAGL, BNR and BAM received a scholarship from CNPq and CAPES.

## Author Contributions Statement

PHACJ designed and supervised the experiments. DSF, GAGL and BRN performed and analyzed experiments and prepared the manuscript. APM and GAGL contributed to the statistical analysis of experimental findings. All authors participated in the discussion of data, and revised the final manuscript.

## Conflict of interest Statement

The authors declare that there are no known conflicts of interest associated with this publication and there has been no significant financial support for this work that could have influenced its outcome.

